# *R*^2^s for Correlated Data: Phylogenetic Models, LMMs, and GLMMs

**DOI:** 10.1101/144170

**Authors:** Anthony R. Ives

## Abstract

Many researchers want to report an *R*^2^ to measure the variance explained by a model. When the model includes correlation among data, such as phylogenetic models and mixed models, defining an *R*^2^ faces two conceptual problems. (i) It is unclear how to measure the variance explained by predictor (independent) variables when the model contains covariances. (ii) Researchers may want the *R*^2^ to include the variance explained by the covariances by asking questions such as “How much of the variance is explained by phylogeny?” Here, I investigate three *R*^2^s for phylogenetic and mixed models. A least-squares *R*^2^_*ls*_ is an extension of the ordinary least-squares *R*^2^ that weights residuals by variances and covariances estimated by the model; it is closely related to *R*^2^_*glmm*_ proposed by Nakagawa & Schielzeth (2013). The conditional expectation *R*^2^_*ce*_ is based on “predicting” each residual from the remaining residuals of the fitted model. The likelihood ratio *R*^2^_*lr*_ was first used by Cragg & Uhler (1970) for logistic regression, and here is used with the standardization proposed by Nagelkerke (1991). These three *R*^2^s are formulated as partial *R*^2^s, making it possible to compare the contributions of mean components (regression coefficients in phylogenetic models and fixed effects in mixed models) and variance components (phylogenetic correlations and random effects) to the fit of models. The properties of the *R*^2^s for phylogenetic models were assessed using simulations for continuous and binary response data (phylogenetic generalized least squares and phylogenetic logistic regression). Because the *R*^2^s are designed broadly for any model for correlated data, the *R*^2^s were also compared for LMMs and GLMMs. *R*^2^_*ls*_, *R*^2^_*ce*_, and *R*^2^_*lr*_ all have good performance, and each has advantages and disadvantages for different applications. These *R*^2^s are computed in the R package rr2 (https://github.com/arives/rr2). [Binomial regression, coefficient of determination, non-independent residuals, phylogenetic model, pseudo-likelihood]

## INTRODUCTION

Researchers often want to calculate a coefficient of determination, an *R*^2^, to give a measure of the amount of variance in their data explained by a statistical model. For ordinary least-squares models (OLS), such as regression and ANOVA, the *R*^2^ is simple to calculate and interpret. Many types of models, however, assume that the errors among response variables are correlated. Phylogenetic generalized least squares models (PGLS) allow the possibility of phylogenetically related species being more similar to each other, leading to phylogenetic correlations in the errors. PGLS models are structurally similar to linear mixed models (LMMs) that include random effects to account for correlations in the residual variation; for example, LMMs can account for correlation between residuals of experimental replicates within the same block. The situation is more complex for models for discrete response variables, such as phylogenetic logistic regression models (PLOG) and generalized linear mixed models (GLMMs). For models of discrete distributions, even perfectly fitting models have residual variation due to the discreteness of the data, and this complicates the interpretation of an *R*^2^.

Correlated errors in statistical models cause two issues for defining an *R*^2^. The first involves assessing the goodness-of-fit of predictor variables (fixed effects) in terms of the explained variance. For standard OLS models, the errors are assumed to be identical and independently distributed, and therefore the variance in the residuals can be calculated directly to give the total variance that is not explained by the model. In models for correlated data, however, the errors are not independently distributed. Therefore, to calculate the “unexplained variance” given by the residuals, it is necessary to deal with the covariances among errors; applying the OLS *R*^2^ to estimates from a model with covariances among errors gives values that are bounded below by -∞ rather than zero (Judge *et al*. 1985 p. 32).

The second issue for defining an *R*^2^ involves assessing the goodness-of-fit of the covariances (random effects) estimated in the model. For phylogenetic models, this is embodied by the question “How much of the data is explained by phylogeny?” The difficulty is that a phylogenetic model can be used to estimate the strength of phylogenetic signal (covariances) in the errors, but the phylogenetic signal does not directly lead to predictions of the fitted data. Desdevises *et al*. (2003) propose an *R*^2^ in which a phylogeny is decomposed into principle components, and the principle components are used as predictor variables in a set of regressions; however, it is not clear how the resulting *R*^2^ maps back to the statistical fit of a model and hence the statistical confidence in its results. An *R*^2^ should ideally be tightly coupled to the fitted model and use this fit to quantify how much of the data are “explained” by the phylogeny.

Here, I assess three *R*^2^s for models that specify non-zero covariances among errors. Although the definitions of *R*^2^s are broad enough to encompass any model specifying an error covariance matrix, I will focus on application to phylogenetic models for continuous (PGLS) and binary (PLOG) data. In addition, I will compare the properties of the *R*^2^s applied to phylogenetic and mixed models, which makes it possible to explore the *R*^2^s in detail. This comparison also validates the *R*^2^s as viable measures of goodness-of-fit for to a broad class of models. This is important, because *R*^2^s should make it possible to assess and compare as wide a range of models as possible (Kvalseth 1985).

The general form of the investigated models is

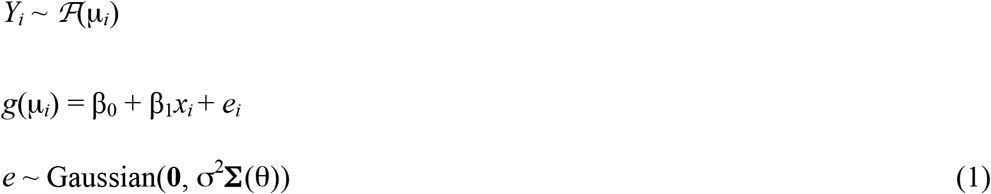

where data *Y_i_* (*i* = 1, …, *n*) are distributed by a member *ℱ* of the exponential family of distributions (McCullagh & Nelder 1989). The parameter µ_*i*_ of distribution *ℱ* is itself a random variable, and applying the link function *g*() to µ_*i*_ gives a linear equation in terms of the predictor variable *x_i_* and an error term *e_i_*. The error term *e_i_* has a multivariate Gaussian distribution with means 0 and covariance matrix σ^2^**Σ**(θ) that may depend on a parameter θ. When the link function *g*() is the identify function, then equation (1) becomes a linear model (e.g., PGLS or LMM). PLOG can be modeled as a phylogenetic GLMM (PGLMM) in which *Y_i_* has outcomes 0 and 1, and the link function *g*() is logit (Ives & Helmus 2011; Ives & Garland 2014; Hadfield 2015). Note that other approaches to phylogenetic logistic regression (Ives & Garland 2010; Ho & Ane 2014) are not structured as PGLMMs, although calculating one of the three *R*^2^s is still possible. Although equation (1) is written for only a single predictor variable *x_i_* and parameter θ (a single random effect in a GLMM), all of the results presented below extend in the obvious way to multiple variables *x_i_* and parameters θ.

The covariances in the residual errors are contained within σ^2^**Σ**(θ). In phylogenetic models, the structure of σ^2^**Σ**(θ) is typically generated under a specific model of evolution (Martins & Hansen 1997b; Lavin *et al*. 2008). For example, in a PGLS using Pagel’s λ branch-length transform (Pagel 1997; Housworth, Martins & Lynch 2004), **Σ**(λ) is the sum of two matrices, **Σ**(λ) = λ**Σ**_BM_ + (1-λ)**I**, where **Σ**_BM_ is the covariance matrix derived under the assumption of Brownian motion (BM) evolution (Felsenstein 1985; Grafen 1989), and **I** is the identity matrix. If λ=1, then the covariances between errors given by BM are proportional to the distance between two tips and their most common ancestor on the phylogenetic tree. If λ=0, the errors are uncorrelated, and 0 < λ < 1 gives intermediate levels of phylogenetic signal. The effect of λ on the covariances among errors can be depicted by adding tip branches to the BM phylogenetic tree that are scaled to make up a proportion λ of the base-to-tip distance (Fig. 1).

**Figure 1.**
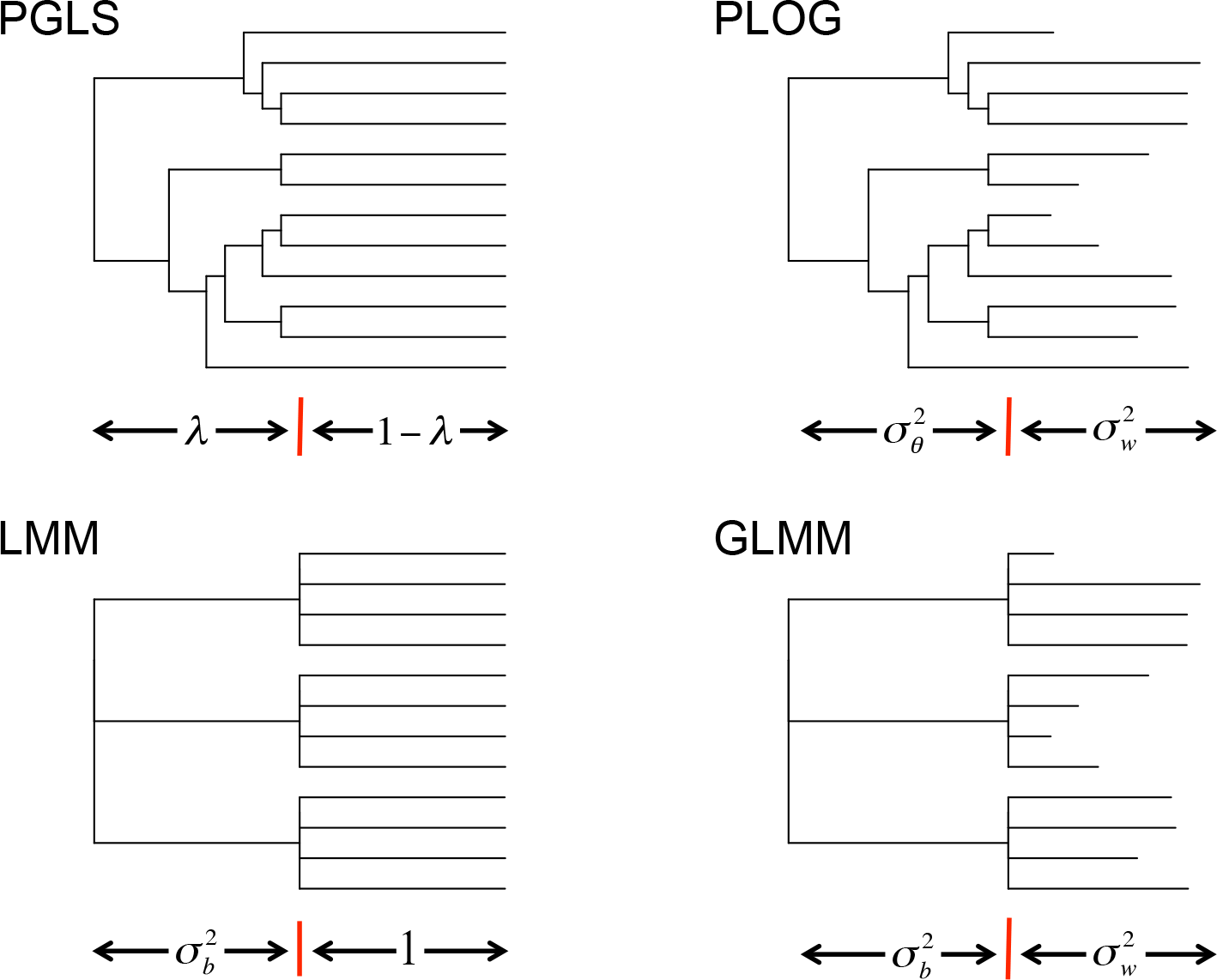
Depictions of the covariance matrices from PGLS, LMM, PLOG, and GLMM. In the PGLS with a Pagel’s λ branch-length transform, the covariance matrix is **Σ**(λ) = λ**Σ**_BM_ + (1-λ)**I**, in which λ determines the strength of phylogenetic signal in the residual errors. In the covariance matrix for LMMs, **Σ**(σ^2^_*b*_) = σ^2^_*b*_ **Σ**_*b*_ + **I**, the variance of the random effect σ^2^_*b*_ is scaled against the variance of the residual errors. For PLOG, phylogenetic signal enters the model as a covariance matrix σ^2^_θ_ **Σ**_BM_, but there is additional variance σ^2^_*w*_ that depends on the difference between observed and predicted values of *Y_i_* which varies for each data point. Similarly, for GLMMs the variance of the random effect is given by **Σ**(σ^2^_*b*_) = σ^2^_*b*_ **Σ**_*b*_, and there is additional variance σ^2^_*w*_ owing to the discreteness of the data.

The similarity between PGLS and LMMs can be seen by depicting the LMM as a tree with branch lengths giving the strength of covariances among errors (Fig. 1). For a model with a single random effect *b*, the covariance matrix is Σ(σ^2^_*b*_) = σ^2^_*b*_ Σ_*b*_ + **I** where Σ_*b*_ is a block-diagonal matrix whose values are 1 for each row *i* and column *j* corresponding to the same level (block) of the random effect (Gelman & Hill 2007). The greater the variance of the random effect σ^2^_*b*_, the greater the covariances among errors within the same level, and the smaller the relative contribution of the residual errors given by the length of the tips of the tree. For comparison with LMMs, GLMMs can also be depicted as a tree, but rather than the residual errors at the tips of the tree having length 1, for models of discrete data the error variance depends on the unavoidable differences between the observation (0 or 1 for binary data) and probability of the observation (taking any value between 0 and 1). The lengths of these variances σ^2^_*w*_ depend on the probability of the observation. I am showing the variances σ^2^_*w*_ only for illustrating the similarities between LMM and GLMM models, and by extension PGLS and PLOG models. In some methods for implementing GLMMs (Schall 1991; Breslow & Clayton 1993), σ^2^_*w*_ (the inverses of the GLM weights; McCullagh & Nelder 1989) are used in the fitting algorithms; in other methods, for example those in the R package ‘lme4’ (Bates *et al*. 2014), they are not used, although they can nonetheless be extracted from fitted models.

The three *R*^2^s presented here partition the “explained” and “unexplained” variances for models with correlated errors such as those depicted in figure 1. Because models can contain multiple parameters, the *R*^2^s compare a full model with a reduced model in which one or more of the parameters are removed; thus, they are partial *R*^2^s that give the explained variance by the components that differ between full and reduced models. The total *R*^2^s are obtained by selecting the reduced model in which there is only an intercept and residuals are independent. Defining partial *R*^2^s has the advantage of being able to ask about the contribution of a single or subset of components to the fit of a model. This makes it possible to exclude coefficients in a model that are not of explicit interest; for example, many phylogenetic models for species traits include body size as one of the predictor variables to factor out body size, and partial *R*^2^s make it possible to assess the goodness-of-fit for the remaining predictor variables. By comparing a model with a phylogeny to a model without, partial *R*^2^s also make it possible to answer the question “How much of the data is explained by phylogeny?"

*R*^2^s can be assessed on multiple grounds (Kvalseth 1985), and here I consider three. First, does the *R*^2^ give a good measure of fit of a model to data? To serve as a basis for assessment, I use the log-likelihood ratio (LLR) of the full and reduced models. The LLR approaches a χ^2^ distribution for large samples and is therefore used for hypothesis tests of full versus reduce models (Judge *et al*. 1985). Also, the LLR is linearly related to the AIC and other measures used for model selection (Burnham & Anderson 2002). Therefore, the LLR is a natural choice to assess *R*^2^s: a good *R*^2^ should be monotonically related to the LLR. Second, can the *R*^2^ separate the contribution of different components of the model to the overall model fit? For the simple case of equation (1) in which there is only a single regression coefficient (β_1_) and a single variance parameter (θ), I ask whether the *R*^2^s can distinguish between the two in their contributions to the fit of the model. Third, does the *R*^2^ give similar values when applied to data generated by the same statistical process? If the values of *R*^2^ when applied to data generated from the same statistical process are all similar, then the *R*^2^ gives a precise measure of goodness-of-fit.

## MATERIALS AND METHODS

There is an extensive literature on *R*^2^s for GLMs and LMMs, and a growing literature for GLMMs (Buse 1973; Cameron & Windmeijer 1996; Cameron & Windmeijer 1997; Kenward & Roger 1997; Menard 2000; Xu 2003; Kramer 2005; Edwards *et al*. 2008; Liu, Zheng & Shen 2008; Orelien & Edwards 2008; Nakagawa & Schielzeth 2013; Jaeger *et al*. 2017), and this literature forms the basis for the *R*^2^s that can be applied to phylogenetic models. The three *R*^2^s take three different approaches to defining “explained variance", the same general approaches considered for LMMs by Xu (2003). *R*^2^_*ls*_ is based on the variance of the residuals in a way that explicitly incorporates their covariances. For models with discrete data, *R*^2^_*ls*_ is defined to closely match *R*^2^_*glmm*_ presented by Nakagawa & Schielzeth (2013). *R*^2^_*ce*_ is based on the difference between the observed data and model predictions (Kvalseth 1985), where the predictions use information from the covariances among errors. *R*^2^_*lr*_ is based upon the information that is gained by adding parameters (regression parameters or covariance terms); it uses the likelihood ratio of full to reduced models as was first proposed for logistic regression (Cragg & Uhler 1970; Maddala 1983; Cox & Snell 1989). For ordinary linear models without correlated errors, these three *R*^2^s are identical to the OLS *R*^2^, but they differ for models with correlated errors.

Here I give a heuristic explanation for the *R*^2^s, and Appendix 1 gives details about their implementation. The starting point in the derivations of the three *R*^2^s is the standard *R*^2^ for continuous data (Buse 1973; Judge *et al*. 1985),

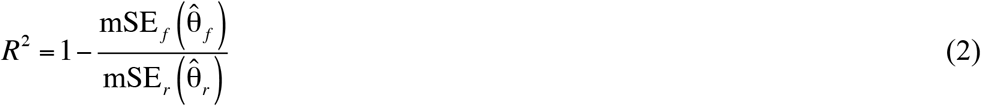

where mSE_*f*_ is the mean squared errors for the full model, and mSE_*r*_ is for the reduced model. For the unadjusted *R*^2^, the mSEs are the mean SEs without correcting for degrees of freedom, so I have used the abbreviation mSE rather than the normal MSE, the mean squared errors corrected for degrees of freedom. Both full and reduced models may contain parameters in vectors θ_*f*_ and θ_*r*_ that involve the variances and covariances among samples which are estimated when the model is fit; for calculating the mSE, these parameters are assumed to be fixed. For a generalized least-squares model (Judge *et al*. 1985),

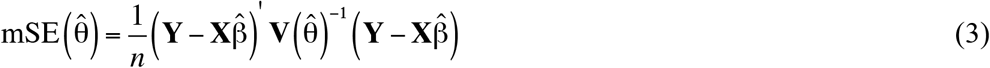

where **Y** is the *n* × 1 vector of response values *Y_i_*, **X** is the *n* × *p* matrix for *p* predictor variables (including the intercept), β̂ is the 1 × *p* vector of estimated regression coefficients (fixed effects), and 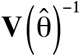 is the inverse of the *n* × *n* matrix **V**(θ) = σ^2^**Σ**(θ) that contains the variances and covariances of the errors. The mSE for OLS models is the special case in which **Σ**(θ) = **I**, which gives the standard *R*^2^.

The key issue in defining *R*^2^s is the scaling of **V**(θ) = σ^2^**Σ**(θ). Setting **V**(θ) = **Σ**(θ) in equation (3), the mSE gives the maximum likelihood estimate of the variance term σ^2^ from equation (1). However, **Σ**(θ) can be rescaled by a constant without changing the fit of the statistical model; the only effect of multiplying **Σ**(θ) by a constant is to change the value of σ^2^ by 1/constant. This would not be an issue if the scaling were the same for full and reduced models, because the scaling would cancel out when dividing mSE_*f*_ by mSE_*r*_. However, it will generally be the case that 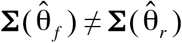; for example, even for LMMs that include the same random effects, 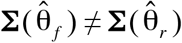 if removing fixed effects from the full model changes the estimated variances of the random effects in the reduce model. Therefore, the scaling is not removed by dividing mSE_*f*_ by mSE_*r*_. Because the scaling determines the estimate of σ^2^, it affects the *R*^2^. The three *R*^2^s presented below differ in how they address the issue of scaling **Σ**(θ).

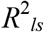

*R*^2^_*ls*_ (for least squares) extends equation (2) in a way that closely matches *R*^2^s that have been proposed for LMMs and GLMMs (Buse 1973; Xu 2003; Edwards *et al*. 2008; Nakagawa & Schielzeth 2013; Jaeger *et al*. 2017). For LMMs, a natural scaling of **Σ**(θ) is to let **Σ**(θ) = **I** + **G**(θ) where **G**(θ) is the block-diagonal matrix containing the variances of the random effects divided by the residual variance (i.e., *σ*^2^_*b*_ in Fig. 1). With this scaling, the residual variance σ^2^ estimated from the LMM equals exactly mSE from equation (3). For the total *R*^2^, the reduced model contains only the intercept and therefore equals the variance of the data. This means that the total *R*^2^_*ls*_ for LMMs differs slightly from *R*^2^_*glmm(c)*_ (Nakagawa & Schielzeth 2013) in which the variance in the data is estimated from the full model, rather than the actual variance of the data. Nonetheless, the values of *R*^2^_*ls*_ and *R*^2^_*glmm(c)*_ will be very close.

For phylogenetic models, scaling **Σ**(θ) is more complicated. For some common branch-length transforms in PGLS that are used to measure the strength of phylogenetic signal, such as the OU transform (Martins & Hansen 1997a; Blomberg, Garland & Ives 2003), the covariance matrix **Σ**(θ) does not separate additively to give terms for the explained versus unexplained variance. Even though Pagel’s λ branch-length transform, **Σ**(λ) = λ**Σ**_BM_ + (1-λ)**I**, does break down into the sum of phylogenetic and non-phylogenetic terms, the non-phylogenetic term (1-λ)**I** cannot be interpreted as the unexplained variance. This is because, for many data sets, the estimate of λ will be 1, which is the expectation under the assumption of BM evolution. If this occurs, this would force an *R*^2^ treating (1-λ)**I** as the unexplained variance to be 1, regardless of the explanatory power of the predictor variances (fixed effects); this property would make the *R*^2^ uninformative. To solve this problem, I propose scaling **Σ**(θ) so that the total branch lengths equal 1; this is equivalent to assuming that the total amount of independent evolution is the same. For a fitted tree with strong phylogenetic signal, scaling **Σ**(θ) to have the total branch lengths equal to 1 will make the base-to-tip distances greater than a fitted tree with no phylogenetic signal. Because this scaling increases the diagonal elements in **Σ**(θ) for greater phylogenetic signal, it will reduce the estimates of σ^2^ and decrease the variance in the residuals that is unexplained by the model. Although this is only a convention (as opposed to a scaling derived from theory), the resulting *R*^2^_*ls*_ performs for PGLS models in a similar way as it does for LMMs.

For discrete models, it is necessary to account for the variation introduced by discrete data (Fig. 1). While there are different ways to do this (Appendix 1), an approach that makes *R*^2^_*ls*_ conform to *R*^2^_*glmm(c)*_ for GLMMs (Nakagawa & Schielzeth 2013) is to replace mSE in equation (2) with 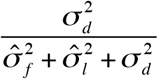 where 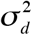 gives the distribution-specific variance attributed to the discreteness of the data; 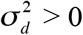 because the predictions from the model are probabilities whereas the observations are discrete values (0 or 1 for binary data). For binary data 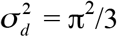. The variance that 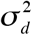 measures is at the level of the transformed parameter value (i.e., *g*(µ_*i*_) in equation (1)) for which the errors are normally distributed. To scale the variance 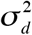 by the variances of the data in the transformed space of *g*(µ_*i*_), it is necessary to divide 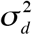 by the total variance that is given by the regression coefficients (fixed effects), 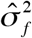, and the errors, 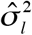. For GLMMs 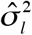 is the estimate of the variance of the random effects. For PGLMMs 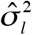 is the estimate of the variance in the Gaussian error term σ^2^**Σ** in which **Σ** is scaled to have base-to-tip branch lengths of 1. Note that in contrast to *R*^2^_*glmm(c)*_ (Nakagawa & Schielzeth 2013), *R*^2^_*ls*_ is explicitly defined as a partial *R*^2^; this not only makes it possible to assess the contributions of different components of the model to goodness-of-fit, it also simplifies application and interpretation of *R*^2^_*ls*_ to LMMs and GLMMs with complex random effects, such as random slope effects (Johnson 2014).

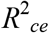

*R*^2^_*ce*_ (for conditional expectation) is based on the variance in the difference between observed and predicted data, 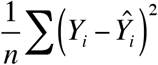. This approach conceptually comes the closest to answer the question of how much variation in the data is explained by the covariances in the model. For the case of LMMs, the predicted values *Ŷ* can be taken as the sum of the fixed effects and the estimate of the value of the random effects. For the LMM in figure 1, this corresponds to the estimated value at the polytomy formed at the node shared by all observations within the same level of the random effect. As the number of observations within each level of the random effect increases, *R*^2^_*ce*_ for LMMs converges to *R*^2^_*ls*_ because the estimates of the values of the random effects become more precise.

As it does for *R*^2^_*ls*_, PGLS poses a complication for *R*^2^_*ce*_: what is the predicted value *Ŷ_i_* ? To parallel *R*^2^_*ce*_ for LMMs, *Ŷ_i_* could be taken as the estimated value at the node immediate below the tip on the phylogeny containing *Y_i_*. For phylogenies with some short terminal branch lengths, however, the estimates for the node underneath *Y_i_* will be determined largely by the value of *Y_i_* itself, leading to very high (and uninformative) *R*^2^s. Therefore, for PGLS I defined *R*^2^_*ce*_ using the estimates *Ŷ_i_* computed by removing the point *Y_i_* from the data set and then estimating *Ŷ_i_* from the predictor variables and the remaining data points. Specifically for equation (1), the expected value of residual *R_i_* = *Y_i_* – (β_0_ + β_1_*x_i_*) from the remaining residuals **R**_[-*i*]_ is

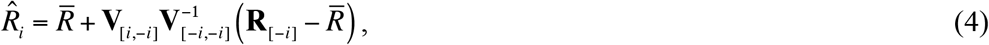

where *R̅* is the GLS mean of the residuals, **V**_[*i,-i*]_ is row *i* of **V** with column *i* removed, and **V**_[-*i,-i*]_ is **V** with row *i* and column *i* removed (Petersen & Pedersen 2012). The predicted value of *Y_i_* is then *Ŷ* = *β*_0_ + *β*_1_*x_i_* + *R̂_i_*. Note that this procedure (removing *Y_i_* before predicting *Ŷ_i_*) could be used for LMMs (and other models), although to make *R*^2^_*ce*_ conform most closely to the structure of LMMs, the LMM *R*^2^_*ce*_ makes predictions *Ŷ_i_* while keeping *Y_i_* in.

The same approach as used for LMMs can by used for GLMMs and PGLMMs. In these cases, the variances are calculated for untransformed values of *Y_i_*, rather than in the Gaussian space of *g*(µ_*i*_) as was done for *R*^2^_*ls*_. These predicted values are given from the estimation algorithms for GLMMs by glmer in the R package lme4 (Bates *et al*. 2014) and for PGLMMs by binaryPGLMM (Ives & Helmus 2011; Ives & Garland 2014) in the R package ape (Paradis & Paradis 2012). Because GLMMs and PGLMMs include the unavoidable variance in *Y_i_* – *Ŷ_i_* due to the discreteness of the data, the estimates *Ŷ_i_* correspond to those values a distance *σ*^2^_*w*_ from the tips of the tree in figure 1.

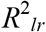

*R*^2^_*lr*_ (for likelihood ratio) is the application of an *R*^2^ proposed for logistic regression (Cragg & Uhler 1970; Maddala 1983; Cox & Snell 1989) and generalized by Magee (1990) and Nagelkerke (1991), and used for LMMs by Kramer (2005). *R*^2^_*lr*_ computed for a range of models in the MuMIn package of R (Barton 2016). For LMMs, *R*^2^_*lr*_ differs from *R*^2^_*ls*_ only in the scaling of **V**(θ). If **V**(θ) is scaled so that the determinant det(**V**(θ)) = 1, then the maximum log likelihood is

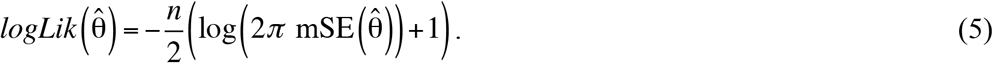

Substituting into equation (2) then leads to

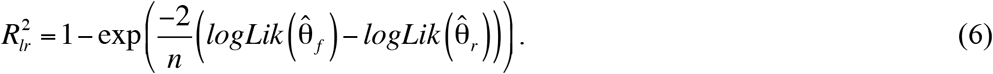

This definition of *R*^2^_*lr*_ in terms of likelihoods extends immediately to any model fit by maximum likelihood estimation, including PGLS and GLMM. However, for discrete data equation (6) does not have a maximum of 1, because the maximum attainable log-likelihood for discrete data is zero. Therefore, Nagelkerke (1991) and Cameron and Windmeijer (1997) proposed dividing by the maximum attainable value, which is equation (6) with 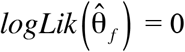; throughout, I have used this Nagelkerke standardization.

The algorithm used by binaryPGLMM to fit equation (1) for binary phylogenetic data uses quasi-likelihoods and does not give a true maximum likelihood that could be used to compute *R*^2^_*lr*_. Therefore, for PLOG models I used phyloglm in the R package phyloglm (Ho & Ane 2014), fitting the model with penalized ML but then using the provided ML values to calculate *R*^2^_*lr*_.

### Simulations for assessment

The simulations to assess the statistical properties of the *R*^2^s applied to LMM, PGLS, GLMM, and PLOG all follow the same strategy. For each, data were simulated using equation (1) for the case with variation in a predictor variable and no covariances (β_1_ > 0, θ = 0), only covariances (β_1_ = 0, θ > 0), or both (β_1_ > 0, θ > 0). For each case, the model parameters were the same for all simulations, so that variation in values of a given *R*^2^ among datasets is caused by random sampling from the same statistical process.

For LMM, data were simulated with the model

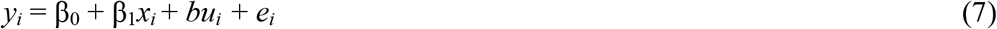

where *x_i_* follows a Gaussian distribution with mean 0 and variance 1, and the random effect *u_i_* has 10 levels, with *b* following a normal distribution with mean 0 and variance θ. I selected parameter values to generate moderate *R*^2^ values. For GLMMs, values from equation (7) without the residual error term *e_i_* were used through a logit link function (equation (1)) to produce binomial probabilities for a binary model. Models were fit using lmer and glmer in the lme4 package of R (Bates *et al*. 2014).

For the PGLS model, to obtain the covariance matrix **Σ**(θ) in equation (1), I first simulated random phylogenetic trees using the rtree function of the ape package of R (Paradis, Claude & Strimmer 2004). Thus, a different tree was simulated for each dataset. The strength of phylogenetic signal was varied using Pagel’s λ transformation which served as the variance parameter θ. Values of *x_i_* were simulated under the BM assumption using the rTraitCont function (Paradis, Claude & Strimmer 2004). The simulated data were fit using penalized maximum likelihood with the function phylolm assuming a Pagel’s lambda transformation in the package phyloglm in R (Ho & Ane 2014). The PLOG model was similar to the PGLS model, but in contrast the predictor variable *x_i_* was assumed to be independently distributed; including phylogenetic signal in *x_i_* caused challenges for model fitting for some simulated datasets, making the simulation studies difficult. Phylogenetic signal in the residuals *e_i_* was controlled by setting **Σ**(θ) = θ**Σ**_BM_ so that in the absence of phylogenetic signal (θ = 0) the simulations conformed to simple logistic regression. To simulate binary data, a logit link function was used in equation (1).

### Simulated examples

I illustrate the three *R*^2^s for two simulated examples. The first involves a PGLS model for two predictor variables and computes the partial *R*^2^s for one of them. For example, suppose a researcher has data on sprint speed in lizards, and the predictor variables are hind leg length and body size (Bauwens *et al*. 1995). Hind leg length is the variable of interest, while body size is a covariate, and therefore the interesting *R*^2^ compares the models with and without hind leg length. I simulated the case of 30 species under BM evolution and compared the cases in which hind leg length (as a proportion of body size) either did or did not show phylogenetic signal. Specifically, the model was

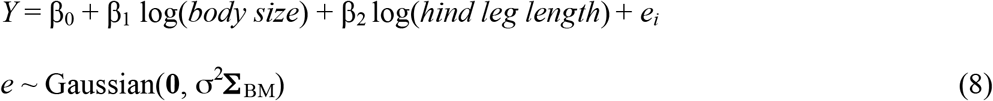

where log(*body size*) was selected from a Gaussian distribution with covariance matrix **Σ**_BM_, and log(*hind leg length*) was selected from a Gaussian distribution with covariance matrix either **I** or **Σ**_BM_. The data were fit using phylolm with Pagel’s λ transformation.

As a second example, I simulated LMMs and binary GLMMs using equation (7), and fit them not only as LMM and GLMMs, but also as PGLS and PLOG models. The fitting as phylogenetic models was performed by converting the covariance matrix given by the random effects in the LMM and GLMM into a phylogenetic tree (Fig. 1) using the vcv2phylo function in ape (Paradis & Paradis 2012). This simulation allows a direct comparison between the *R*^2^s applied to mixed versus phylogenetic models.

## RESULTS

The *R*^2^s were assessed according to the three properties: (i) their ability to measure goodness-of-fit as benchmarked by the LLR of full model and the model with only an intercept, (ii) whether they can partition sources of variation in the model as partial *R*^2^s, and (iii) how precise is their inference about goodness-of-fit. Property (iii) treats the *R*^2^s as if they were estimators of goodness-of-fit and asks how variable are the estimates when applied to repeated simulations from the same model (e.g.,; Cameron & Windmeijer 1996). A more comprehensive assessment is given in Appendix 2.

### Goodness-of-fit

Figure 2 plots the total *R*^2^s against the corresponding LLR. All *R*^2^s were positively related to the LLR, which is a minimum requirement for an *R*^2^. *R*^2^_*lr*_ shows a monotonic relationship with LLR, which is necessarily the case due to the definition of *R*^2^_*lr*_ (equations (5), (6)). For the remaining *R*^2^s, values for a given LLR were generally lower for simulations in which variation was produced only by the fixed effect (β_1_ > 0, θ = 0; Fig. 1, blue circles). This implies that, relative to the LLR, these *R*^2^s were attributing less “explained” variance to regression coefficients (fixed effects) than covariances parameters (phylogeny and random effects).

**Figure 2.**
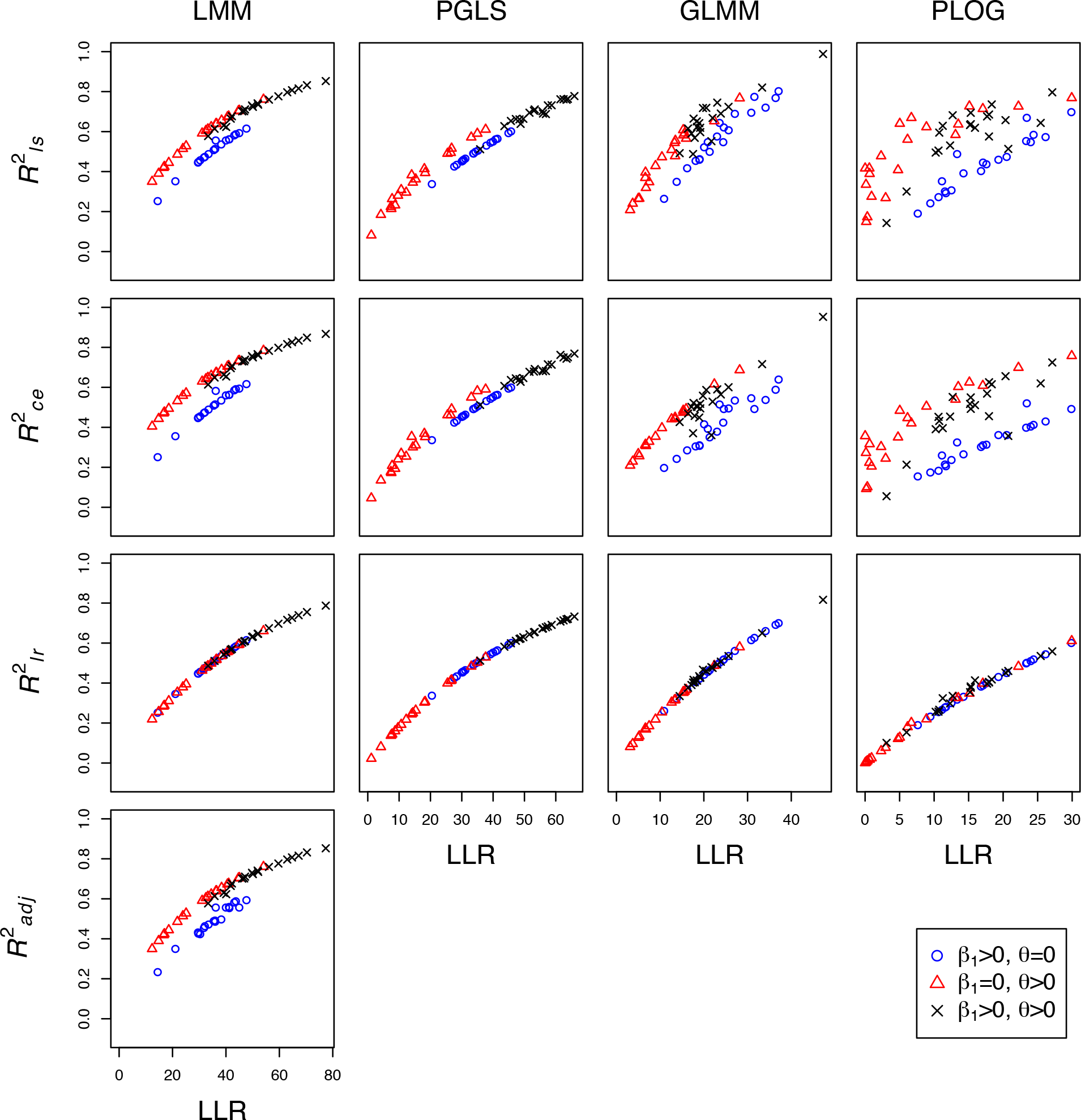
Results for LMM, PGLS, GLMM, and PLOG simulations giving *R*^2^_*ls*_, *R*^2^_*ce*_, *R*^2^_*lr*_, and the OLS *R*^2^_*adj*_ versus the log likelihood ratio (LLR) between full model and reduced model containing only an intercept. All simulated data had 100 samples. For LMM, the simulation model (equation (7)) contained a fixed effect with β_1_ = 0 or 1, and a random effect *u_i_* with 10 levels and variance θ = 0 or 1.5. The binomial (binary) GLMM was similar but with β_1_ = 0 or 1.8, and θ = 0 or 1.8. For PGLS, β_1_ = 0 or 1, and the strength of phylogenetic signal θ = λ = 0 or 0.5; for PLOG β_1_ = 0 or 1.5, and θ = 0 or 2. The LMM was fit using lmer (Bates *et al*. 2014); the GLMM was fit using glmer (Bates *et al*. 2014); the PGLS was fit using phylolm (Ho & Ane 2014); and for PLOG LLR and *R*^2^_*lr*_ were fit using phyloglm (Ho & Ane 2014), and *R*^2^_*ls*_ and *R*^2^_*ce*_ were fit using binaryPGLMM (Ives & Garland 2014). For reduced models without variance parameters, fitting was done using lm and glm.

For the LMM, I included the adjusted *R*^2^, *R*^2^_*adj*_ computed from OLS regression by treating the random effect as a categorical fixed effect. *R*^2^_*ls*_ and *R*^2^_*adj*_ were almost identical. This correspondence implies that *R*^2^_*ls*_ gives an *R*^2^ that is interpretable in the same way as the standard *R*^2^_*adj*_ but generalized to LMMs.

All of the *R*^2^s other than *R*^2^_*lr*_ showed greater scatter in their relationships with LLR for the simulations of binary data (GLMM and PLOG). In part, this is due to the difficulty of estimating variance parameters θ in binomial models. For example, there is more scatter in *R*^2^_*lr*_ for GLMM simulations than LMM simulations, even though the criterion for calculating the *R*^2^_*s*_ (the log likelihoods) are the same. The scatter seems particularly large for *R*^2^_*ls*_ and *R*^2^_*ce*_ applied to PLOG simulations, although this case requires some technical discussion. For PLOG, the LLR was obtained from phyloglm using penalized maximum likelihood, whereas *R*^2^_*ls*_ and *R*^2^_*ce*_ were estimated from the model fit by binaryPGLMM using the pseudo-likelihood. The phyloglm estimate of phylogenetic signal, λ, tended to absorb at zero even when the estimate λ from binaryPGLMM was positive; therefore, *R*^2^_*ls*_ and *R*^2^_*ce*_ could be positive even when the LLR was zero. Previous comparison between phyloglm and binaryPGLMM showed that they have similar performances but do not necessarily give the same conclusions about the presence of phylogenetic signal for the same dataset (Ives & Garland 2014).

### Partitioning sources of variation

The partial *R*^2^_*ls*_, *R*^2^_*ce*_, and *R*^2^_*lr*_ were generally able to partition sources of variation between components of a model, in particular between regression coefficients (fixed effects) and covariance parameters (random effects). Simulations with β_1_ > 0 and θ = 0 should have partial *R*^2^s for β_1_ that are positive and partial *R*^2^s for θ that are zero (blue circles, Fig. 2). Simulations with β_1_ = 0 and θ > 0 should have partial *R*^2^s for β_1_ that are zero and partial *R*^2^s for θ that are positive (red triangles, Fig. 2). Simulations with β_1_ > 0 and θ > 0 should have both partial *R*^2^s > 0 (black x’s, Fig. 2). Because the values of β_1_ and θ were the same whether or not the other was zero, the partial *R*^2^s for β_1_ should be the same for simulations with θ = 0 (blue circles) as for simulations with θ > 0 (black x′s), and the partial *R*^2^s for θ should similarly be the same for β_1_ = 0 (red triangles) and β_1_ > 0. For continuous data (LMM and PGLS), all three *R*^2^s had similar performance and similar values of the partial *R*^2^s (see also Appendix 2). For binary data (GLMM and PLOG), the three *R*^2^s showed more scatter, which in large part is due to the greater statistical challenge of estimating regression coefficients and variance parameters from discrete data. This is seen, for example, in the GLMM and PLOG simulations with β_1_ > 0 and θ > 0 which sometimes gave a partial *R*^2^_*lr*_ for θ of zero (black x’s); these cases occur when the estimate of θ was zero even though a non-zero value was used in the simulations.

### Inference about underlying process

The ability of *R*^2^s to infer the fit of the statistical process to the model depends on the precision of the estimates of *R*^2^. Figure 4 plots the mean values of the *R*^2^s with 66% and 95% inclusion intervals for simulated datasets with sample sizes 40, 60, …, 160. For LMMs and GLMMs, there were 10 levels of the random effect; datasets were produced by first simulating 160 samples (16 replicates at each level) and then randomly removing two replicates at each level to reduce the sample size in steps of 20. For PGLS and PLOG, each dataset at each sample size was simulated independently.

For LMM simulations, *R*^2^_*ls*_, *R*^2^_*ce*_, and *R*^2^_*adj*_ showed similar patterns (Fig. 4), reflecting the fact that they give very similar values (Fig. 2, Appendix 2). Mean values did not change with sample size, and there was only moderate increase in variability among simulations with decreasing sample size. In contrast, mean values of *R*^2^_*lr*_ decreased with decreasing sample size. This probably reflects the information that is lost when estimating the model parameters. In contrast to LMM simulations, the PGLS simulations showed less change in the means of *R*^2^_*lr*_ and *R*^2^_*ce*_ with sample size, presumably because there were more covariances among samples (i.e., the covariance matrix had more non-zero elements) than in the LMM with few replicates per level.

For the GLMM, both *R*^2^_*ls*_ and *R*^2^_*ce*_ had somewhat higher variances (less precision) than *R*^2^_*lr*_. The greater variation in values of *R*^2^_*ls*_ and *R*^2^_*ce*_ compared to *R*^2^_*lr*_ may occur because *R*^2^_*ls*_ and *R*^2^_*ce*_ depend on estimates from the models (random effects for *R*^2^_*ls*_ and fitted values for *R*^2^_*ce*_) whereas *R*^2^_*lr*_ depends on likelihoods. Thus, *R*^2^_*ls*_ and *R*^2^_*ce*_ are compromised when the estimates are poor, as is particularly the case when sample sizes are small. For PLOG, the variances in *R*^2^_*ls*_ and *R*^2^_*ce*_ were similar to *R*^2^_*lr*_ (Fig. 4). This is likely because estimates of phylogenetic signal (λ = θ) were well-bounded, in contrast to the variance in random effects in the GLMMs.

### Simulated examples

In the simulated example of sprint speed regressed on log body mass (*x*_1_) and log hind limb length (*x*_2_) (equation (8)), whether or not log hind limb length showed phylogenetic signal had a large effect on the partial *R*^2^s for the effect of log hind limb length (Table 1). In the fitted PGLS models for both datasets, the parameter estimates and log likelihoods were similar, and the only indication that phylogenetic signal in hind limb length affected the fit of the model was the *p*-value for the regression coefficient for hind limb length (*P* = 0.012 when *x*_2_ had phylogenetic signal and *P* << 0.001 when it did not). The partial *R*^2^_*ls*_, *R*^2^_*ce*_, and *R*^2^_*lr*_ for hind leg length were 0.21-0.23 when hind leg length had phylogenetic signal, and 0.71-0.89 when it did not. In contrast, the partial *R*^2^s for phylogenetic signal (reduced model with λ = 0) and the total *R*^2^s (reduced model with *x*_1_ = *x*_2_ = λ =0) did not differ much between simulations. The partial *R*^2^s for hind limb length depended upon the phylogenetic signal in hind limb length because when there is phylogenetic signal and hind limb length is removed from the model, much of the information is recaptured in the phylogenetic signal of the residual variation. This example illustrates the value of having partial *R*^2^s that can assess the role of predictor variables separately from other variables like body size that are not of specific interest.

**Table 1:** Simulation example of sprint speed regressed on log body mass (*x*_1_) and log hind limb length (*x*_2_). For the simulation, the regression coefficients for log(*body size*) and log(*hind leg length*) were β_1_ = 1 and β_2_ = 0.5, and the intercept was β_0_ = 0 (equation (8)); residual variation was given by BM evolution, so λ = 1. Hind limb length was simulated under BM evolution (left table) or as a normal random variable (right table), in both cases with variance 1.

| coefficient | estimate | $P$ |
| --- | --- | --- |
| intercept | -0.46 | 0.50 |
| $x_1$ | 0.79 | 0.006 |
| $x_2$ | 0.55 | 0.012 |
| $\lambda$ | 1 | |

$N = 30$ , logLik = -17.76
| reduce model | $R^2_{ls}$ | $R^2_{ce}$ | $R^2_{lr}$ |
| --- | --- | --- | --- |
| $x_2=0$ | 0.21 | 0.23 | 0.21 |
| $\lambda=0$ | 0.69 | 0.89 | 0.72 |
| $x_1=x_2=\lambda=0$ | 0.79 | 0.93 | 0.81 |

No phylogenetic signal in $x_2$
| coefficient | estimate | $P$ |
| --- | --- | --- |
| intercept | -0.49 | 0.46 |
| $x_1$ | 0.75 | 0.008 |
| $x_2$ | 0.55 | <<0.001 |
| $\lambda$ | 1 | |

$N = 30$ , logLik = -17.12
| reduced model | $R^2_{ls}$ | $R^2_{ce}$ | $R^2_{lr}$ |
| --- | --- | --- | --- |
| $x_2=0$ | 0.71 | 0.89 | 0.79 |
| $\lambda=0$ | 0.71 | 0.90 | 0.74 |
| $x_1=x_2=\lambda=0$ | 0.88 | 0.96 | 0.90 |

In the second example (Table 2), a LMM and a GLMM were simulated and then the data were fit using LMM and GLMM, and also PGLS and PLOG by converting the covariance matrix given by the random effects into a phylogeny (Fig. 1). *R*^2^_*ls*_, *R*^2^_*lr*_, and *R*^2^_*ce*_ were computed for the total model, as well as partial *R*^2^_*s*_ for the fixed effect x and the random effect θ. As expected, *R*^2^_*lr*_ was the same for mixed and phylogenetic models. All values for *R*^2^_*ce*_ were also close between mixed and phylogenetic models. For the LMM simulation, *R*^2^_*ls*_ calculated for the LMM model was very close to both *R*^2^_*ce*_ and the *R*^2^_*ols*_ computed by treating the random effect as a categorical fixed effect. However, the values of *R*^2^_*ls*_ calculated from PGLS were lower. This occurred because the scaling of the covariance matrix Σ for LMM and PGLS is different; for LMM *R*^2^_*ls*_ scales Σ so that the residual error corresponds to 1, whereas for PGLS *R*^2^_*ls*_ scales Σ so that the total branch lengths equal 1. Although this gives different values of *R*^2^_*ls*_ for LMM and PGLS, it avoids forcing *R*^2^_*ls*_ to equal 1 when the residual error matches its expectation under BM evolution. For the GLMM simulation, *R*^2^_*ls*_ from the fitted GLMM is higher than *R*^2^_*ls*_ from the fitted PLOG, although *R*^2^_*ls*_ for both models are higher than *R*^2^_*lr*_ and *R*^2^_*ce*_.

**Table 2:** Illustrative simulation comparing *R*^2^s for data simulated from a LMM and fitted with LMM and PGLS, and for data simulated from a binary GLMM and fitted with GLMM and PLOG. Data were simulated from equation (7) with 10 levels of 10 observations each (*n* = 100), a random effect with variance 2, a normally distributed *x* with mean 0 and variance 1, and for the LMM a residual error variance of 0.5. PGLS and PLOG were fit by converting the covariance matrix given by the random effects into a phylogeny (Fig. 1). Columns labeled “Mixed” and “Phylo” correspond to LMM and GLMM, and PGLS and PLOG, respectively. For the LMM, the adjusted *R*^2^_*adj*_ was computed from the OLS model fix by treating the random effect as a fixed effect with 10 levels.

| | | $R^2_{lr}$ | | $R^2_{ce}$ | | $R^2_{ls}$ | | $R^2_{adj}$ |
| --- | --- | --- | --- | --- | --- | --- | --- | --- |
|  |  | Mixed | Phylo | Mixed | Phylo | Mixed | Phylo |  |
| LMM |  |  |  |  |  |  |  |  |
|  | total | 0.916 | 0.916 | 0.955 | 0.945 | 0.950 | 0.853 | 0.955 |
| | partial $x$ | 0.722 | 0.722 | 0.764 | 0.763 | 0.764 | 0.490 | 0.764 |
| | partial $\theta$ | 0.913 | 0.913 | 0.954 | 0.943 | 0.949 | 0.848 | 0.954 |
| GLMM |  |  |  |  |  |  |  |  |
|  | total | 0.317 | 0.317 | 0.471 | 0.468 | 0.591 | 0.550 |  |
| | partial $x$ | 0.141 | 0.141 | 0.161 | 0.159 | 0.343 | 0.297 | |
| | partial $\theta$ | 0.298 | 0.298 | 0.461 | 0.459 | 0.577 | 0.535 | |

## DISCUSSION

*R*^2^_*ls*_, *R*^2^_*ce*_, and *R*^2^_*lr*_ are presented here with focus on phylogenetic models, although they are broadly applicable to models with correlated errors. Below, I first address their specific application to phylogenetic models using mixed models as a reference, and then give general recommendations.

### Applications to LMM, PGLS, GLMM, and PLOG

For both continuous data (LMM and PGLS) and discrete data (GLMM and PLOG), all *R*^2^s had good performance. For the simple model with a single regression coefficient β_1_ and single covariance parameter θ (equation (7)), all *R*^2^s were reasonable measures of goodness-of-fit, as assessed against the log likelihood ratio between full and reduced models (Fig. 2). Nonetheless, *R*^2^_*ls*_ and *R*^2^_*ce*_ gave lower partial *R*^2^ values for the regression coefficient β_1_ relative to the partial *R*^2^ values for the covariance parameter θ in comparison to *R*^2^_*lr*_ and the log likelihood ratio, LLR (Fig. 2). Also, although all three *R*^2^s gave very similar values for the same dataset with continuous data, the values differed more for discrete data fit with either GLMM or PLOG (Fig. 2 and Appendix 2). This is reflected in general by the decreased precision of the *R*^2^s applied to discrete data, as measured by the variation in values when fit to data simulated under the same parameter values (Fig. 4). All *R*^2^s were capable of identifying whether β_1_ or θ was responsible for the fit of the model to the data as determined by the partial *R*^2^s; when β_1_ or θ were zero in the simulations, the partial *R*^2^s for β_1_ or θ, respectively, were low (Fig. 3). However, the partial *R*^2^s for GLMM and PLOG tended to be more variable and less conclusive than for LMM and PGLS (Fig. 3). Finally, *R*^2^_*lr*_ decreased as sample sizes decreased especially for LMMs but also for GLMMs (Fig. 4). This is an understandable consequence of the loss of information to separate full and reduced models when there are fewer data. Nonetheless, it is an undesirable property, just as the change in the unadjusted OLS *R*^2^ is undesirable.

**Figure 3.**
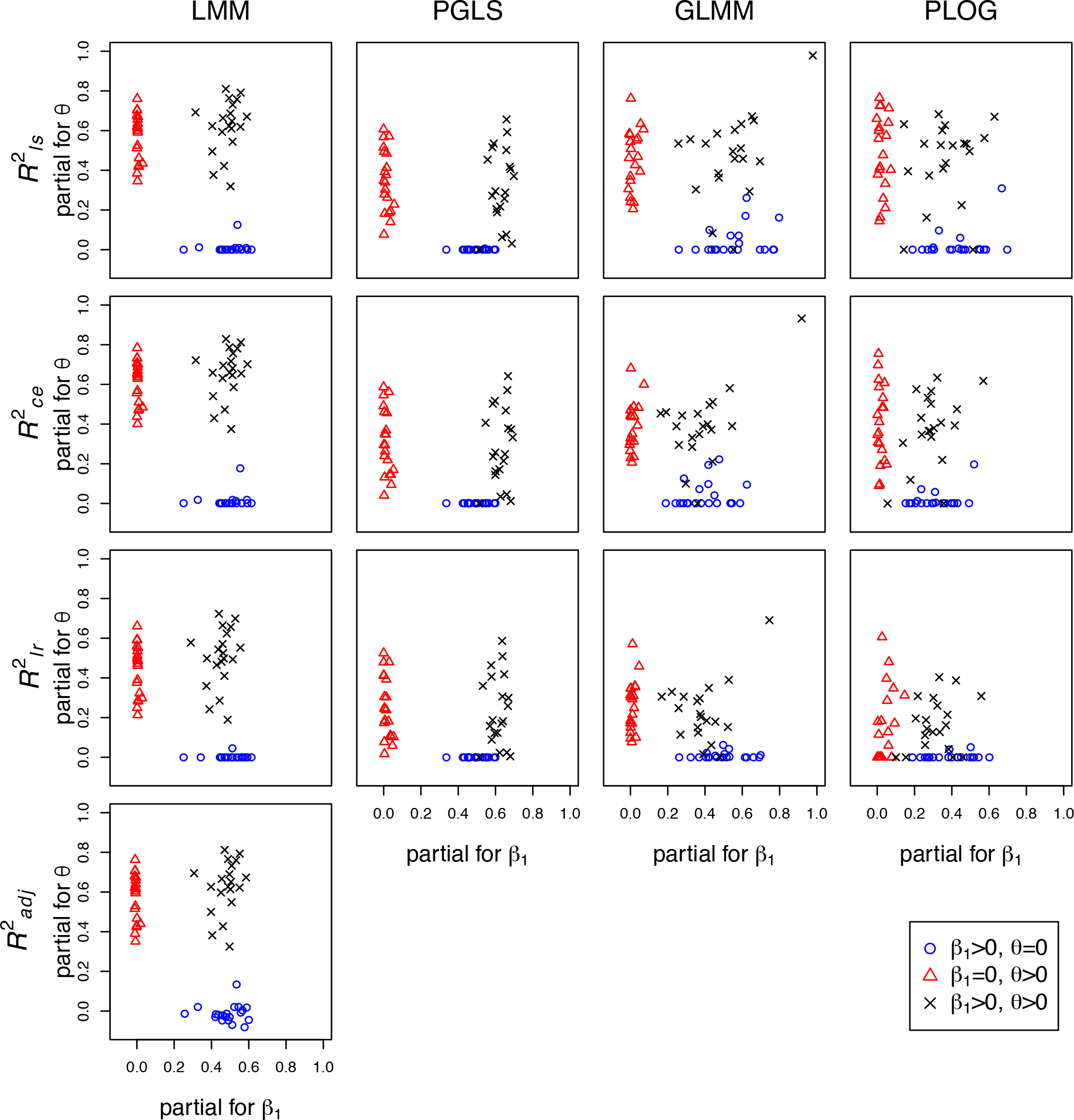
Results for LMM, PGLS, GLMM, and PLOG simulations giving partial values of *R*^2^_*ls*_, *R*^2^_*ce*_, *R*^2^_*lr*_, and *R*^2^_*adj*_. The partial *R*^2^ for β_1_ was calculated using the reduced model in which θ is removed, and for the partial *R*^2^ for θ the reduced model had β_1_ removed. The simulated data and fitting methods are the same as in figure 1.

**Figure 4.**
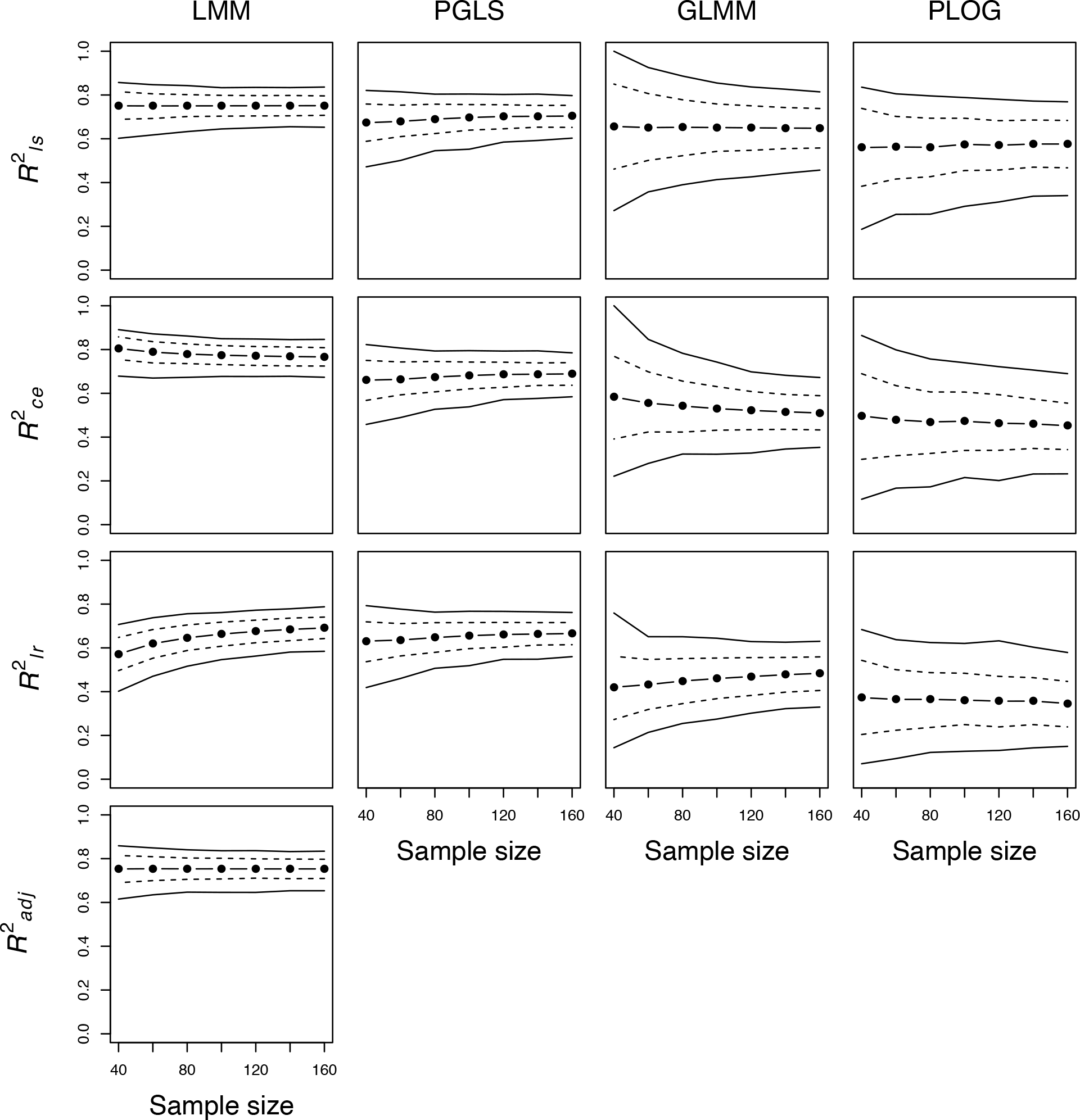
Results for LMM, PGLS, GLMM, and PLOG simulations showing means, 66% and 95% inclusion intervals for *R*^2^_*ls*_, *R*^2^_*ce*_, *R*^2^_*lr*_, and *R*^2^_*adj*_ versus sample size. For all simulations 1000 data sets were analyzed at each sample size. Parameter values were: LMM, β_1_ = 1, θ = 1.5; PGLS, β_1_ = 1, θ = 0.5; GLMM, β_1_ = 1.8, θ = 1.8; and PLOG, β_1_ = 1.5, θ = 2.

The poorer performance of all three *R*^2^s for GLMM and PLOG relative to LMM and PGLS in terms of partitioning sources of variation (Fig. 3) and precision (Fig. 4) is due to the greater challenges of fitting discrete data. This will affect the *R*^2^s differently if they are differently sensitive to the fitting. *R*^2^_*ls*_ is calculated from fitted variances in a model; *R*^2^_*ce*_ is calculated from the fitted values of *Y_i_*; and *R*^2^_*lr*_ is calculated from the likelihood. Therefore, the three *R*^2^s will be sensitive to the precision with which each of these attributes is estimated. For GLMMs, the precision of *R*^2^_*lr*_ was slightly greater than the other two *R*^2^s, although this did not appear to be the case for PLOG.

Although it is hard to argue in favor of one *R*^2^ over the others on the basis of their performance in the simulations, *R*^2^_*ls*_ has the advantage of producing *R*^2^_*glmm(c)*_ (Nakagawa & Schielzeth 2013) as a special case when applied to LMMs and GLMMs. Furthermore, because *R*^2^_*ls*_ (like the other *R*^2^s) is defined as a partial *R*^2^, it makes application to LMMs and GLMMs more flexible and general, allowing subsets of fixed and random effects to be analyzed, and also more-complex structures like random slopes (Johnson 2014). Nonetheless, a partial *R*^2^_*glmm*(*c*)_ comparable to *R*^2^_*ls*_ can easily be defined as

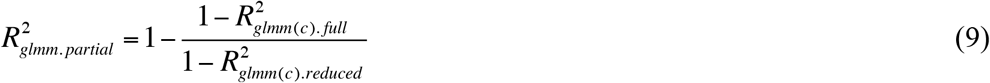

Although the relationship between *R*^2^_*ls*_ and R^2^_*glmm(c)*_ might argue in favor of *R*^2^_*ls*_ over *R*^2^_*ce*_ and *R*^2^_*lr*_, when applied to data simulated from LMM and GLMM, *R*^2^_*ls*_ values calculated from fitted LMM and GLMM were different from the values calculated from PGLS and PLOG fitted to the same data (Table 2). This highlights a weakness of *R*^2^_*ls*_: a decision has to be made about how to scale the covariance matrix **Σ** depending on the fitted model, and the resulting values of *R*^2^_*ls*_ will depend on this decision.

### Recommendations

An ideal *R*^2^ would make it possible to compare among different models and among different methods used to fit the same model (Kvalseth 1985 properties of a good R2 #4 and #5). *R*^2^_*ls*_ and *R*^2^_*ce*_ can be used for any model and fitting method that estimates the covariance matrix (*R*^2^_*ls*_) and/or fitted values (*R*^2^_*ce*_); for example, they could be used to compare LMMs fit with ML vs. REML, or binary phylogenetic models fit with ML (e.g., phyloglm; Ho & Ane 2014) or quasi-likelihood (e.g., binaryPGLMM; Ives & Garland 2014). Nonetheless, *R*^2^_*ls*_ and *R*^2^_*ce*_ have a disadvantage in terms of generality. For *R*^2^_*ls*_ a decision must be made about how to scale the covariance matrix **V**(θ) (equation (3)), and for *R*^2^_*ce*_ a decision has to be made about how values of *Y_i_* are predicted. The conventions I used for LMMs and PGLS differed. In contrast, *R*^2^_*lr*_ is restricted to models that are fit with ML estimation; however, if ML is used for fitting, then values of *R*^2^_*lr*_ can be compared across different types of models. This applies to any type of data and model fit with ML estimation.

An ideal *R*^2^ should also be intuitive (Kvalseth 1985 property #1). However, intuitive is in the eye of the beholder. *R*^2^_*ls*_ is the most similar to the OLS *R*^2^, which grounds *R*^2^_*ls*_ in the familiar and intuitive OLS framework. *R*^2^_*ce*_ predicts the data from covariances estimated in the model, and therefore could be viewed as the most intuitive way to relate the variance explained by regression coefficients (fixed effects) to that explained by variance parameters (random effects). *R*^2^_*lr*_ is also related to the OLS *R*^2^: in LMMs and PGLS, *R*^2^_*lr*_ only differs from *R*^2^_*ls*_ by the way in which the covariance matrix **V**(θ) (equation (3)) is scaled, and this provides a link between *R*^2^_*lr*_ and the OLS *R*^2^ through *R*^2^_*ls*_. This said, however, I suspect that different researchers would rank the intuitiveness of *R*^2^_*ls*_, *R*^2^_*ce*_, and *R*^2^_*lr*_ differently.

*R*^2^s are often used as “summary statistics" to describe the fit of a model to data in a way that does not involve statistical inference about the underlying stochastic process that generated the data: “How does the model fit these data?" rather than “How much does the model infer about the process that generated the data?" Should *R*^2^s be judged as a summary statistic? I think not. All the *R*^2^s showed high variation among simulations of the same model with the same parameters, especially when sample sizes were small (Fig. 4). This means that how the model fits a specific dataset involves a lot of chance, and hence one should not get too excited about a high *R*^2^, or too discouraged about a low one. *R*^2^s are best treated as inferential statistics, that is, as functions of a data-generating process that are themselves random variables (Cameron & Windmeijer 1996; Nakagawa & Schielzeth 2013). As an inferential statistic, *R*^2^_*lr*_ most directly ties to hypothesis testing between full and reduce models using a likelihood ratio test. For me, this tips the balance to favor *R*^2^_*lr*_ over the others.

## ACKNOWLEDGMENTS

I thank Ted Garland, Daijiang Li, Shinishi Nakagawa, Eric Pedersen, and Joe Phillips for wonderfully insightful comments that helped to clarify this article.

## SUPPLEMENTARY MATERIAL

Appendix 1: Details about the implementation of the *R*^2^s.

Appendix 2: More comparisons among the *R*^2^s in the R package rr2. supplementary files: source R scripts for computing *R*^2^_*ls*_, *R*^2^_*lr*,_ and *R*^2^_*ce*_ with examples.

